# Knock-in of labeled proteins into 5’UTR enables highly efficient generation of stable cell lines

**DOI:** 10.1101/2020.08.03.234252

**Authors:** Faryal Ijaz, Koji Ikegami

## Abstract

Stable cell lines and animal models expressing tagged proteins are important tools for studying behaviors of cells and molecules. Several molecular biological technologies have been applied with varying degrees of success and efficiencies to establish cell lines expressing tagged proteins. Here we applied CRISPR/Cas9 for the knock-in of tagged proteins into the 5’UTR of the endogenous gene loci. With this 5’UTR-targeting knock-in strategy, stable cell lines expressing Arl13b-Venus, Reep6-HA, and EGFP-alpha-tubulin were established with high knock-in efficiencies ranging from 50 to 80%. The localization of the knock-in proteins were identical to that of the endogenous proteins in wild-type cells and showed homogenous expression. Moreover, the expression of knock-in EGFP-alpha-tubulin from the endogenous promoter was stable over long-term culture. We further demonstrated that the fluorescent signals were enough for a long time time-lapse imaging. The fluorescent signals were distinctly visible during the whole duration of the time-lapse imaging and showed specific subcellular localizations. Altogether, our strategy demonstrates that 5’UTR is a ‘hotspot’ for targeted insertion of gene sequences and allows the stable expression of tagged proteins from endogenous loci in mammalian cells.

## Introduction

Labeling proteins of interest with fluorescent proteins or small tags is one of the most popular methods in life science to investigate the localization or/and roles of the proteins. Especially, the technique is powerful and essential when reliable antibodies against the proteins of interest are not available. The simplest way is the transient overexpression of the labeled proteins of interest by transfecting cells with plasmid vectors harboring promoters, such as CMV promoter. Some researchers generate stable cell lines that express the labeled proteins from the expression unit randomly and occasionally inserted into the genome. However, both the transient and stable expression of the proteins of interest under exogenous promoters has some disadvantages; causing artifacts or toxicity due to the overexpression or/and silencing of gene expression during long time maintenance of the generated stable cell lines (Chen et al., 2011, Domenger and Grimm, 2019, Xiong et al., 2019, Shin et al 2020).

Insertion of tagged genes of interest (GOI) into the genomic safe harbor (GSH) loci, such as ROSA26 allows the new genes to be expressed without disrupting the gene function (Friedrich and Sorano, 1991). So far, several fluorescent reporters or tagged GOIs, e.g. fusion proteins, have been introduced into the ROSA26 locus (Irion et al., 2007, Tchorz et al., 2012, Li et al., 2017, Imayoshi et al., 2020). Traditionally, a transgene of interest flanked by splice acceptor sequence and a stop cassette is inserted into an Xba1 restriction site within the first intron of the ROSA26 gene (Zambrowicz et al., 1991). Exogenous promoters, e.g. CMV promoter, are introduced to drive high transgene expression in some cases where the moderate strength of ROSA26 promoter does not achieve a desirable level of transgene expression (Mao et al., 1999, Srinivas et al., 2001, Muzumdar et al., 2007, Madisen et al., 2010). More recently, CRISPR/Cas9 system has also been utilized to generate ROSA26 knock-ins (KI) by designing gRNA targeting ROSA26 locus (Chu et al., 2016, Ma et al., 2017, Wu et al., 2018). ROSA26 locus is still prone to the heterogeneity of the protein expression and has issues regarding the long-term stability of transgene overexpression in stable cell lines while it is amenable for KI of GOI, though (Chen et al., 2011, Shin et al 2020).

Among the genome editing technologies, the CRISPR/Cas9 system offers the greatest ease of use (Cong et al. 2013, Mali et al., 2013) and have been optimized for targeted gene insertion of peptide tags, reporter genes and genes of interest at endogenous gene loci through a variety of repair pathways. Homology-dependent recombination (HDR) uses a homologous repair template to accurately repair the double-strand breaks (DSBs) (Capecchi, 1989, Takata et al., 1998). The system has been utilized for the targeted integration of transgenes. V5 or Flag epitope sequences, for example, have been inserted at the 3’end of endogenous transcription factor loci of DNA or RNA-binding proteins in HEK293T, HepG2, and MCF7 cells (Savic et al., 2015, Nostrand et al., 2017). Recently, HDR-USR (universal surrogate reporter) system has been utilized to improve the KI efficiency of EGFP integration cassettes into the GAPDH locus of HEK293T cells (Yan et al., 2019). However, accomplishing high-efficiency KI in mammalian cells by HDR can be challenging and frequently involves artificial manipulation of cellular DNA repair systems because the rate of HDR-mediated DNA repair from DSBs is naturally extremely low in mammalian cells. (Chu et al., 2015, Lee et al., 2018, Maruyama et al., 2015). We also struggled to acquire a positives clone for the KI mIMCD-3 cell line expressing IFT81 tagged with a YNL fluorescent tag and could only get a single positive clone out of 100 candidates through HDR mediated gene editing (Phua et al., 2017).

Contrary to HDR, the non-homologous end-joining (NHEJ) is the predominant mode of repair in the eukaryotic cells (Lieber et al, 2003) and does not need a homologous DNA template, which simplifies genome editing strategies. NHEJ has been successfully used to generate KI in vivo and variety of cell lines (Maresca et al., 2013, Auer et al., 2014, Suzuki et al., 2016, Lackner et al., 2015, Schmidt-Burgk et al., 2016, Sawatsubashi et al., 2018, Manna et al., 2019). Several strategies have been employed to generate NHEJ-dependent CRISPR/Cas9-mediated KIs. In knock-in blunt end ligation (KiBL), PCR amplicons lacking homology arms were introduced at the target site into HEK293 cells by using two sgRNAs simultaneously to facilitate the precise excision of the intervening sequence (Geisinger et al., 2016). Another technique, named homology-independent targeted integration (HITI), enables the targeted gene insertion into non-dividing cells in vitro and in vivo (Suzuki et al., 2016). The sgRNA/spCas9 complex introduces DSBs in the genomic target resulting in two blunt ends. The donor DNA also contains the identical sgRNA/spCas9 sequences in the reverse direction. The linearized donor DNA is inserted at the target site via NHEJ (Suzuki et al., 2016, Suzuki and Belmonte 2018). In another approach, KI of large reporter genes via universal reporter system using sg-A target site taken from prokaryotic DNA sequence allowed the integration of promoter-less ires-eGFP fragment into GAPDH locus of LO2 cells and human embryonic stem cells (He et al., 2016). In one strategy, ‘self-cleaving’ GFP-plasmids containing universal gRNAs were used to generate NIH/3T3 and N2a stable cell lines (Tálas et al., 2017). Manna and colleagues demonstrated the use of CHoP-In (CRISPR-mediated, Homology-independent, and PCR-product Integration) approach to make HeLa, HEK293T, and NRK stable cell lines (Manna et al. 2019). Still, the efficiency of targeted insertion of fluorescent proteins or small tags at the endogenous loci of GOI through NHEJ remains low in a wide variety of mammalian cell lines due to the occurrence of frameshift during NHEJ-dependent repair (Bachu et al., 2015, Schmidt-Burgk et al., 2016).

Here, we describe a highly efficient CRISPR/Cas9 mediated homology-independent KI of tagged GOI into the 5’ untranslated region (5’UTR) of endogenous gene loci in mammalian cells. We show that our method is fast and can efficiently drive the stable and homogenous expression of labeled proteins from endogenous loci.

## Materials and Methods

### Cas9 and sgRNA constructs

Broad Institutes GPP sgRNA Designer was used to design the sgRNAs (https://portals.broadinstitute.org/gpp/public/analysis-tools/sgrna-design). CRISPR/Cas9 target sequences (20-bp target and 3-bp PAM (underlined) used in this study are as follows: Mouse Arl13b 5’UTR-targeting (ACGTCAGCACGTCGACGCGG<u>GGG</u>), Mouse Reep6 5’UTR-targeting (AGGTTGCGGGCGTGCTTCAG<u>TGG</u>), Mouse Tub1a5’UTR-targeting (CCTCGCCTCCGCCATCCACC<u>CGG</u>), and Mouse ROSA26 Intron 1-targeting (CGGCACTGGGAATCCCCGTG<u>CGG</u>)

To construct Cas9- and gRNA-expression plasmid vectors, each a 20-bp target sequence was sub-cloned into CMV-Cas9-2A-GFP plasmid backbone (ATUM, CA, USA). To construct donor plasmid vectors, p5’UTRgRNA-Arl13b-Venus, p5’UTRgRNA-Reep6-HA, and p5’UTRgRNA-EGFP-Tub1a, the CMV promoter was removed from the expression vectors of Arl13b-Venus, Reep6-HA and EGFP-alpha-tubulin fusion proteins. Next, the 20-bp target sequence and 3-bp PAM sequence were sub-cloned upstream of the fusion proteins sequences. For pROSA26gRNA-EGFP-Tub1a, 20-bp target sequence and 3-bp PAM sequence were sub-cloned directly upstream of the CMV promoter.

### Cell culture and transfection

NIH/3T3 cells (ATCC CRL-1658) were cultured in DMEM-High Glucose (Wako, Japan) with 10% FBS. mIMCD-3 (mouse inner medullary collecting duct-3) cells (ATCC CRL-2123) were maintained in Dulbecco’s modified Eagle’s medium (DMEM)/Ham’s F-12 (Wako, Japan) supplemented with 10% fetal bovine serum (FBS). All cells were incubated at 37°C with 5% CO_2_. Ciliogenesis in NIH/3T3 was induced by replacing the medium with DMEM-High Glucose with 1% FBS.

Polyethylenimine (PEI) was used for transfection (Longo et al., 2013). Transfection complexes were prepared by mixing PEI and DNA at a ratio of 3:1 in weight. For transient overexpression, expression plasmid vectors of fusion proteins, Arl13b-Venus, Reep6-HA, and EGFP-Tub1a were transfected into NIH/3T3 or mIMCD-3 cells lines. To generate stable cell lines, Cas9/gRNA expression vectors and donor vectors were co-transfected using PEI. The following combinations of Cas9/gRNA expression vectors and donor vectors were used: NIH/3T3 cells expressing Arl13b-Venus (p5’UTRgRNA-Arl13b-Venus / CMV-Cas9-2A-GFP-mArl13b-5’UTRgRNA); mIMCD-3 cells expressing EGFP-Tub1a from endogenous promotor (p5’UTRgRNA-EGFP-Tub1a / CMV-Cas9-2A-GFP-mTub1a-5’UTRgRNA); mIMCD-3 cells expressing Reep6-HA from endogenous promotor (p5’UTRgRNA-Reep6-HA / CMV-Cas9-2A-GFP-mReep6-5’UTRgRNA); and mIMCD-3 cells expressing EGFP-Tub1a from ROSA26 locus (pROSA26gRNA-EGFP-Tub1a / CMV-Cas9-2A-GFP-ROSA26-gRNA).

To compare the stability of expression of fusion proteins from endogenous promoter and CMV promoter, mIMCD-3 cells stably expressing EGFP-Tub1a from endogenous locus without an exogenous promoter or from ROSA26 locus with the CMV promoter were seeded at the density of 2.0×10^5^ on the cover slips in a 12-well plate. After 2 days of incubation, cells were fixed and immuno-stained as described. Samples were collected every week for up to 12 weeks.

### Generation of stable cell lines

The workflow is shown in Fig.1. Cells co-transfected with Cas9/gRNA expression vectors and donor vectors were treated with 1 mg/mL G418 for the selection. During the G418 selection, cells were expanded into larger dish. Then, selected G418-resistant cells were collected and single cell-cloned into 96-well plates with the limiting dilution-culture method at 0.2∼0.3 cell/well. Cells were allowed to grow for 1-2 weeks. For selection of monocolonies, microscopic observations were done to monitor single cell colony formation and confluency. Selected colonies were expanded into the duplicate of multi-well plates from which one culture was used to make frozen stocks for subsequent use and the other culture for screening. Screening of positive clones was done via western blot analyses. G418 was added throughout the cloning and screening steps until the stocks were made from positive clones while decreasing the antibiotic concentration at each step after selection to 0.2-0.5 mg/mL. G418 was omitted when the stable cell lines were maintained for experiments.

### Antibodies

The antibodies used in this study are as follows: alpha-tubulin (mouse mAb DMIA; T9026; Sigma), Arl13b (mouse mAb N295B/66; ab136648; Abcam), Arl13b (rabbit pAb; 17711-1-AP; Protein Tech), GFP (rabbit pAb; 598; MBL), HA.11 Epitope Tag (mouse mAb 16B12; 901502; Biolegend), Alexa fluorophore-conjugated secondary antibodies (Thermo) for immunofluorescence microscopy, and horseradish peroxidase-conjugated secondary antibodies (Jackson Immuno Research Laboratories) for western blot analyses.

### Western blot analyses

Cell lysates were harvested by adding 1x SDS-PAGE sample buffer to the confluent cells and heated at 95°C for 5 minutes and loaded on to an acrylamide gel. Following protein transfer, PVDF membranes were blocked with 5% BSA/TBST for 1 h at room temperature. Next, primary antibodies diluted in 1% BSA/TBST were added and the membranes were incubated overnight at 4°C. Following incubation and washing, blots were incubated with HRP-conjugated secondary antibodies for 1 h at room temperature and developed using ECL prime (GE, UK).

### Immunofluorescence microscopy

Cells were fixed with 4% paraformaldehyde (PFA, pH 7.5) for 30 min at 37°C. Cells were blocked and permeabilized with 5% normal goat serum containing 0.1% Triton X-100 in PBS for 1 h at room temperature. Then, cells were incubated overnight with primary antibodies diluted in 5% normal goat serum/0.1% triton X-100/PBS. Cells were washed with PBS and incubated for 1 h with Alexa Fluor-conjugated secondary antibodies (1:500) and DAPI (1:1000; DOJINDO). After washing, cover glass containing cells were mounted on glass slides with Vecta Shield mounting medium (Vector Laboratories, USA). Images were captured with an epifluorescence microscope (Leica DMI3000B) equipped with a lens (20X NA: 0.70 40X NA: 0.85 or 100X NA: 1.40) and a MiChrome 5 Pro CMOS camera (Tucsen Photonics, China).

### Time-Lapse Imaging

Arl13b-Venus-expressing NIH/3T3 cells were grown to confluence in a 35-mm glass bottom dish (Matsunami, Japan). Cells having a primary cilium were imaged every one minute for total of 1.5 h using a confocal microscope (Olympus FV1000) equipped with an oil immersion lens (60X, NA:1.35). mIMCD-3 cells expressing EGFP-alpha-tubulin were grown to 50% confluency in a 35-mm glass bottom dish (Matsunami, Japan) and were imaged every one minute for total of 1 h using a confocal microscope (Olympus FV3000) equipped with an oil immersion lens (60X, NA: 1.40). Taxol (Wako, Japan) was added at a final concentration of 5 µM to the dish containing KI cells after 5 minutes into imaging.

### Image processing and data analysis

At least 10 images were obtained per sample and fluorescence intensities were measured using ImageJ (https://imagej.nih.gov/ij/). Primary cilium length was measured using “line” tool of ImageJ. GFP-positive cells and DAPI-stained nuclei were counted using “Multi-point” tool of ImageJ. Microsoft Excel was used to calculate average, standard error of the mean (s.e.m.) and statistical significance based on Student’s *t*-test. Probability values <0.05 were considered significant. Graphs were plotted with KyPlot (KyensLab Inc.) and GraphPad Prism.

## Results

### Scheme of CRISPR/Cas9-mediated 5’UTR-targeting knock-in of tagged proteins

We selected a gRNA sequence to guide Cas9 endonuclease to induce DSBs in the 5’UTR of the coding genes in the genomic DNA allowing targeted integration of tagged GOI upstream of the coding sequence and downstream of the endogenous promoters (Fig. 1A). This approach allows the simultaneous disruption of endogenous allele and the expression of tagged protein from the endogenous promoter, replacing the endogenous protein with the tagged protein. We used mammalian expression vectors that had been constructed previously from pcDNA derivatives for the transient transfection and overexpression of proteins of interest as the materials to construct donor vectors. Donor vectors were readily constructed by simply replacing the CMV promoter with the gRNA sequence and PAM sequence that are identical to those in the 5’UTR of target gene (Fig. 1A). The donor vectors constructed from conventional mammalian expression vectors are ready for use as they contained polyadenylation signal sequences to stabilize transcribed RNA and the expression cassette of antibiotic resistance genes (Fig. 1A). Following co-transfection of the donor vector and the Cas9/gRNA-expression vector, the linearized donor vector cleaved by Cas9 endonuclease was integrated at the target site in genomic DNA during the repair of Cas9-mediated DSB through the NHEJ pathway (Katoh et al., 2017), which occurred in the 5’UTR of GOI in our case (Fig. 1A). The tagged GOI expression arise only from the correct insertion at the endogenous GOI loci as it was promoter-less (Fig. 1A). The antibiotic resistance gene was used for the positive selection of cells in which the integration of donor vectors into the genomic DNA occurred (Fig. 1A).

**Figure 1.**
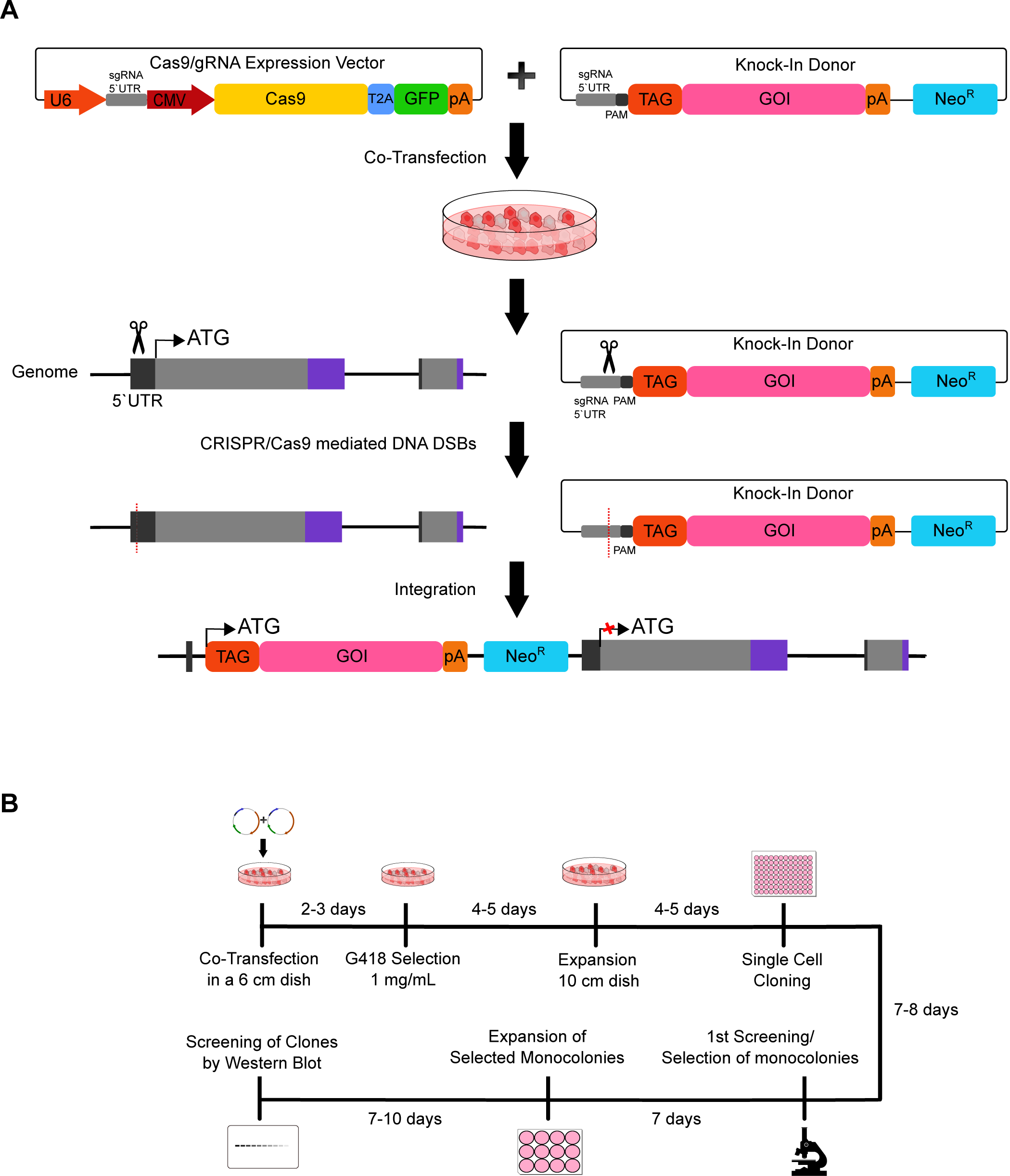
Knock-in of tagged genes of interest into 5’UTR of endogenous gene locus. **(A)** Schematics of the donor plasmid and targeting strategy for CRISPR/Cas9-mediated homology independent knock-in of tagged gene of interest (GOI) at the 5’UTR of the endogenous gene locus. After co-transfection of the KI donor and Cas9/gRNA expression vectors, Cas9 introduce a double-strand break (DSBs; red dotted lines) upstream of the protospacer adjacent motif (PAM) sequence (dark grey) in the 5’UTR of genomic DNA and tagged GOI in the donor plasmid. NHEJ directed repair leads to the integration of the cassette into the genome. (Neo^R^: neomycin resistance gene cassette) **(B)** Timeline for the preparation of stable cell lines. After co-transfection of donor and target vectors, clones were selected with G418 (1mg/mL) and monoclonies were expanded after 14 days of culture. Western blotting was done to screen for positive clones.

We established a workflow to generate stable mammalian cell lines expressing tagged proteins from the endogenous promoters (Fig. 1B). The workflow enables to acquire desired clones for 1-1.5 months (Fig. 1B). It took 7-10 days to finish the selection of G418-resistant transfected and knock-in candidates after starting the culture (Fig. 1B). The cloning step took about 2-3 weeks, depending on cell types; mIMCD-3 cells grow faster while NIH/3T3 cells proliferate slower in our hands (doubling time: 22∼24 h versus 36∼45 h). The screening was completed in 1-2 days as it was a conventional western blot analysis. After finishing the screening, the culture media were free of G418.

### Generation of a NIH/3T3 stable cell line expressing C-terminally Venus-tagged Arl13b, a ciliary marker protein, from endogenous loci for live cell imaging

To test the efficacy of our approach, we first targeted a protein that is accumulated in a highly restricted region in cell. We selected ADP-ribosylation factor-like protein 13B (Arl13b), which is highly accumulated within a primary cilium, a tiny protrusion on a cell surface with several micro meter in length and ∼200 nm in diameter (Caspary et al., 2007). It is widely used as a marker of the primary cilium in studies about cilia (Hua and Ferland, 2017). Mouse Arl13b gene is located on chromosome 16 and consists of 10 exons. The 5’UTR region of this gene is made up of 297 bp. We selected gRNA target sequence located 20-bp downstream from 5’ end of 5’UTR (21-40 bp; Fig. 2A, magenta). Upon transfection, CMV-dependently expressed Cas9 endonuclease introduces a DSB 3-bp upstream of the Cas9 recognition sequence PAM (41-43 bp; Fig. 2A, orange) and in the same sequence in the KI donor vector. During NHEJ repair, the released donor sequence consisting of Arl13b-coding region fused with the Venus at the N-terminus and Neo^R^ cassette integrates at the target site, leading to the expression of Arl13b-Venus protein from the endogenous promoter (Fig. 2A).

**Figure 2.**
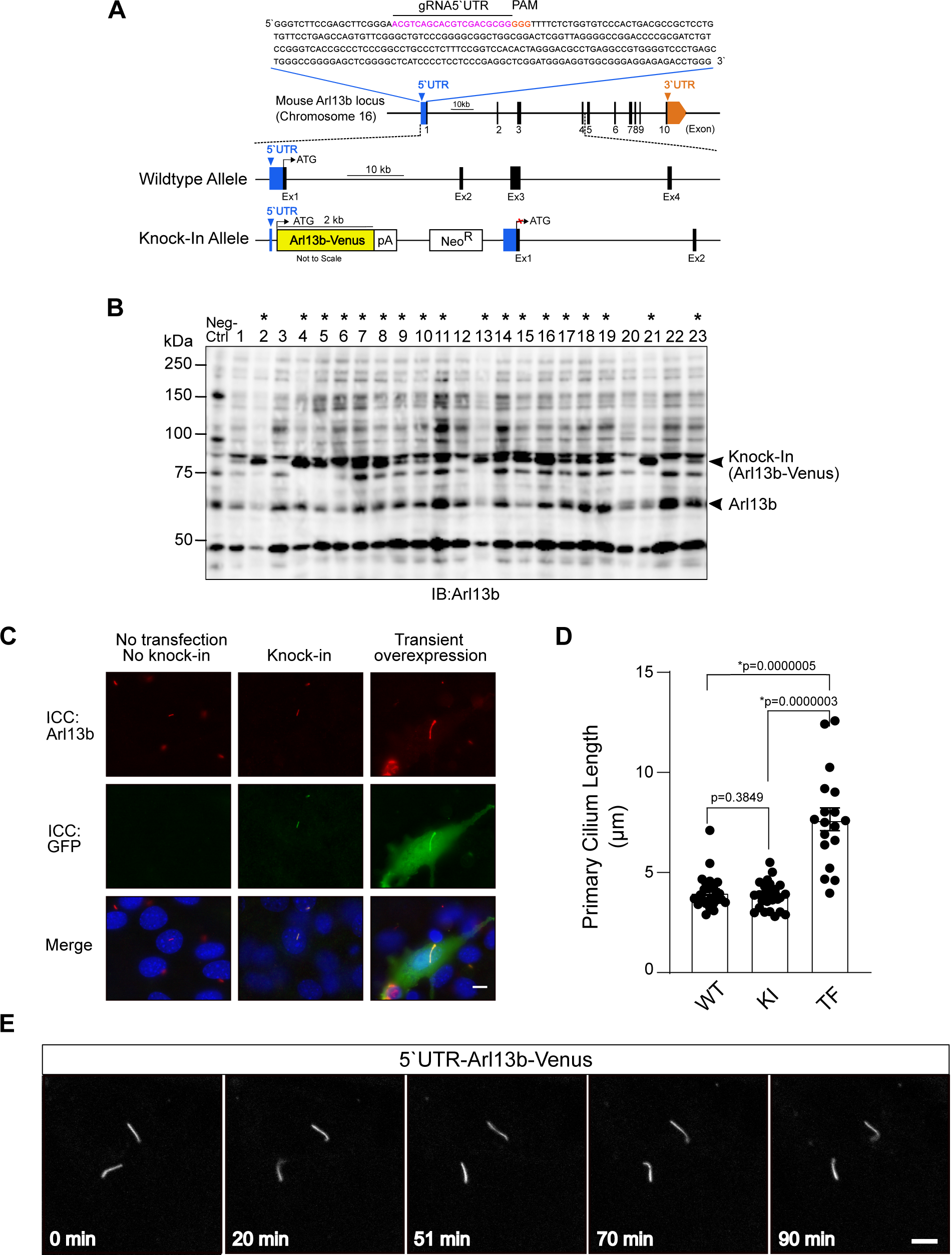
CRISPR/Cas9 mediated knock-in of Arl13b-Venus into the 5’UTR of Arl13b locus in NIH/3T3 cells. **(A)** Diagram of the mouse Arl13b locus. The gRNA-5’UTR target sequence (magenta) upstream of the protospacer adjacent motif (PAM, orange) in the 5 untranslated region (5’UTR) are indicated. **(B)** Screening of Arl13b-Venus positive clones by western blotting. Lysate from wildtype NIH/3T3 cells was used as negative control. Clones positive for integration are marked by asterisks *. **(C)** Fluorescence microscopy images after immunocytochemistry showing cells with primary cilium stained for ciliary marker (Arl13b; red), Arl13b-Venus (GFP; green) and the nucleus (DAPI; blue) in control (no knock-in and no transfection of Arl13b-Venus), knock-in cells and cells with transient overexpression of Arl13b-Venus. Scale bar, 10 μm. **(D)** Quantitative analysis of primary cilium length from C. Data are shown as the mean ± s.e.m. (WT: wildtype; KI: knock-in; TF: transfection; *, *p* < 0.05). **(E)** Time-lapse confocal microscopy images of the primary cilium with endogenous expression of Arl13b-Venus. Time stamps are shown at the bottom (Scale bar, 5 μm).

Applying this strategy, 23 G418-resistant clones were picked and characterized with western blot analysis. The native Arl13b has a molecular weight of 60 kDa (Fig. 2B; lower arrow head), while knock-in Arl13b-Venus bands were detected at 87 kDa (Fig. 2B; upper arrow head). Nineteen of these 23 clones were screened as correctly targeted clones (* of Fig. 2B; 83%). To examine whether our strategy kept the localization of Arl13b, we induced ciliogenesis in NIH/3T3 cells stably expressing Arl13b-Venus and checked the expression of Arl13b-Venus in primary cilium through immunostaining with antibodies against Arl13b and Venus proteins. Venus signals, which were further amplified with immunostaining with anti-GFP antibodies, were specifically detected in primary cilia labeled with anti-Arl13b antibodies in KI cells (Fig. 2C; middle row). Cytoplasm of KI cells were negative for Venus signals and Arl13b signals (Fig. 2C; middle row). The specificity of anti-GFP antibodies immunoreactivity was verified by observing that no GFP staining in wild-type negative controls (Fig. 2C; left row). In cells transiently overexpressing Arl13b-venus upon the CMV promoter activity, both Venus signals and Arl13b signals were remarkably stronger than those of KI cells (Fig. 2C; right row). Markedly, overexpression of Arl13b-Venus resulted in mislocalization, i.e. “leak”, of Arl13b-Venus in cytoplasm (Fig. 2C; right row).

We further examined whether KI of Arl13b-Venus kept the morphology of primary cilia normal. The length of the primary cilium in the KI cells stably expressing Arl13b-Venus was comparable to that of wild-type negative control cells (Fig. 2C, D). In contrast, the length of primary cilium with transient overexpression of Arl13b-Venus was abnormally long, exceeding more than 10 µm in some cases (Fig. 2C, D). These results strongly demonstrate that our approach is capable of keeping localization and function of Arl13b in physiological conditions.

To check whether our approach is suitable for live cell imaging, we performed time-lapse imaging of Arl13b-Venus-positive primary cilia in the KI cell line using a confocal laser scanning microscope. The fluorescent signals of Arl13b-Venus in the primary cilium were readily and clearly detected in a single focal plane, as primary cilia of NIH/3T3 tend to protrude parallel to cell surface or glass surface (Ijaz and Ikegami, 2019). The fluorescent signals of Arl13b-Venus was maintained thoroughly throughout the imaging over the course of 90 min with images taken every 1 min (Fig. 2E; Movie1).

### Generation of an mIMCD-3 stable line expressing a membrane protein tagged with a small tag at C-terminal for detecting the target proteins

We next tried knocking in a membranous protein that is not fused to a fluorescent protein, rather tagged with a small tag, as a case where researchers are difficult to access reliable antibodies to detect the target protein. We chose a short membrane protein, receptor expression-enhancing protein 6 (Reep6), a member of the REEP/Yop1 family of proteins that spans three times plasma membrane and is localized on the ER membrane (Saito et al 2004; Bjork et al., 2013). Mouse Reep6 gene is present on chromosome 10, comprises of 6 exons and its 5’UTR region is 278 bp long. The 20-bp gRNA (233-252 bp; Fig. 3A, magenta) was selected 232-bp downstream from 5’ end of 5’UTR. After co-transfection of Cas9/gRNA-expression and donor vectors, Cas9-induced cleavage in the targeted allele and the donor vector was repaired via the NHEJ-mediated repair pathway allowing targeted integration and expression of Reep6-HA from the endogenous locus (Fig. 3A).

**Figure 3.**
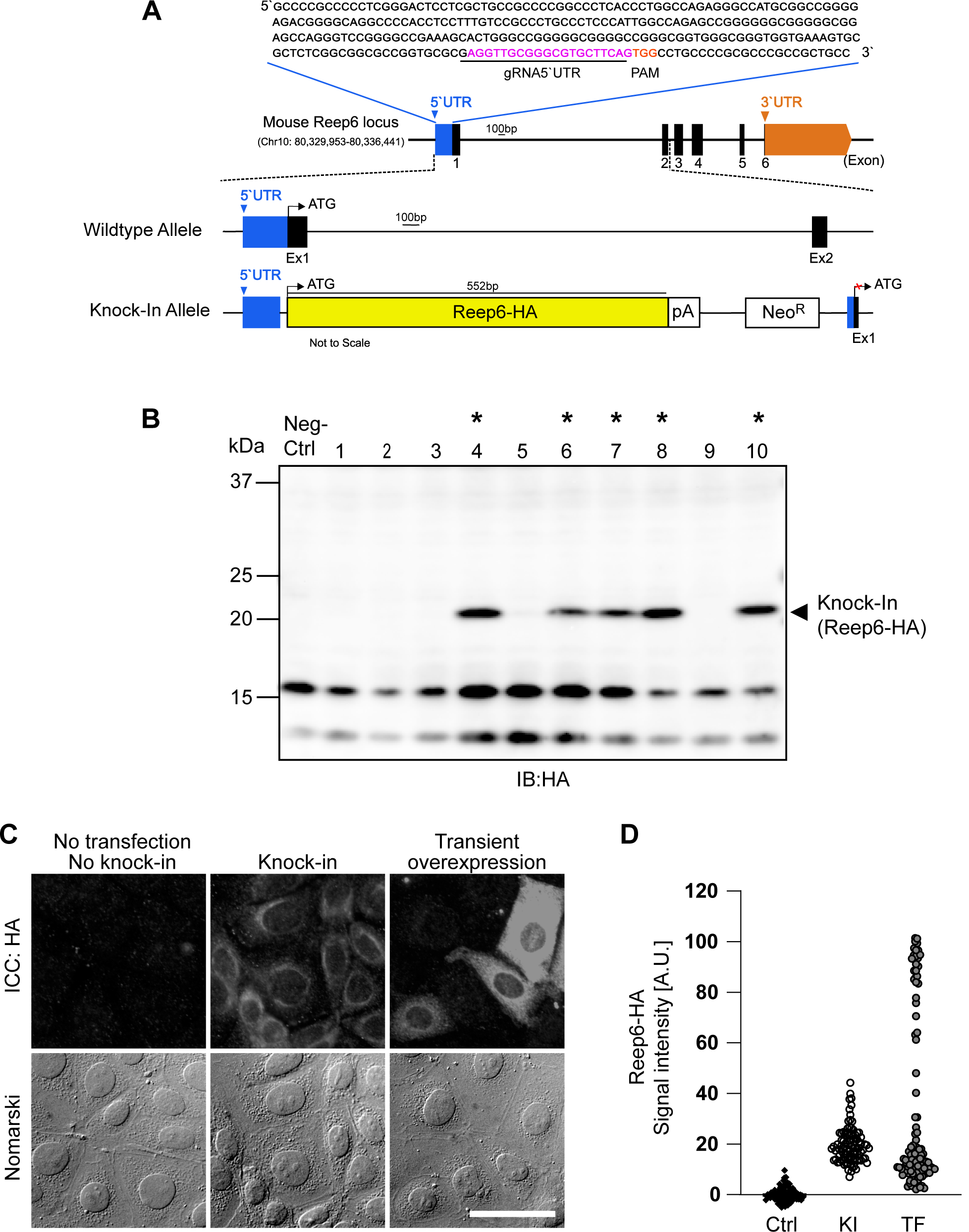
CRISPR/Cas9 mediated knock-in of Reep6-HA into the 5’UTR of Reep6 locus in IMCD3 cells. **(A)** Diagram of the mouse Reep6 locus. The gRNA-5’UTR target sequence (magenta) upstream of the protospacer adjacent motif (PAM, orange) in the 5′untranslated region (5’UTR) are indicated. **(B)** Screening of Reep6-HA positive clones by western blotting. Lysates from wildtype IMCD3 cells was used as negative control. Clones positive for integration are marked by asterisks *. **(C)** Fluorescence microscopy images after immunocytochemistry with anti-HA antibody showing expression of Reep6-HA in knock-in cells, cells with transient overexpression of Reep6-HA and control (no knock-in and no transfection of Reep6-HA). Scale bar, 50 μm. **(D)** Quantification of Reep6-HA levels as described in **C** (ctrl: control; KI: knock-in; TF: transfection).

Five out of ten selected clones (50%) were positive for integration of Reep6-HA as confirmed by using western blot analysis with an anti-HA antibody (Fig. 3A; *). Reep6-HA protein band was detected at 22 kDa (arrow head) and was absent in negative control (Fig. 3B). The anti-HA antibody was used in this study as reliable antibodies against Reep6, which were able to distinguish Reep6-knockout cells from wild-type cells, were not available. The western blot shows that the expression level of knocked in Reep6-HA was enough to be detected with the commercially available and well-distributed antibody for the small tag, HA (Fig. 3B).

To further investigate the localization of knocked-in Reep6-HA, we compared them in immunofluorescent staining. The signals detected with the anti-HA mAb showed the ER-like distribution, which was accumulated surrounding on half a side of nuclei, in the KI cell line (Fig. 3C; middle row). The negative control, i.e. wild-type, cells were not stained with the anti-HA antibody at all (Fig. 3C; left row). Transient overexpression of Reep6-HA upon the CMV promoter by plasmid transfection resulted in extraordinary expression of Reep6-HA in some cells, where the intracellular distribution of Reep6-HA looked impaired with signals diffused thoroughly in cytoplasm (Fig. 3C; right row). In addition, the quantitation of fluorescent intensities demonstrates that the expression level of Reep6-HA was moderate and homogenous in the KI cells while the cells transiently overexpressing Reep6-HA showed heterogeneous expression of Reep6-HA along with extraordinarily enhanced expression (Fig. 3D).

### Generation of an mIMCD-3 stable line expressing N-terminal-EGFP-tagged α-Tubulin housekeeping gene for live cell imaging of responses to treatment

We also generated a stable cell line expressing a fluorescent protein-tagged housekeeping gene. We chose alpha-tubulin as the target housekeeping gene to carry out live cell imaging of microtubule polymerization upon drug treatment. Mouse alpha-tubulin 1A (Tub1a) gene is present on chromosome 15, consists of 4 exons and its 5’UTR region is 250 bp long. The 20 bp gRNA (209-228 bp; Fig. 4A, magenta) was selected 208-bp downstream from 5’ end of 5’UTR. Once the knocking in was achieved, the inserted EGFP-Tubulin was expressed under the direct control of the endogenous promoter (Fig. 4A). Using this strategy, 11/20 (55%) clones were found to express EGFP-Tubulin (Fig. 4B; *).

**Figure 4.**
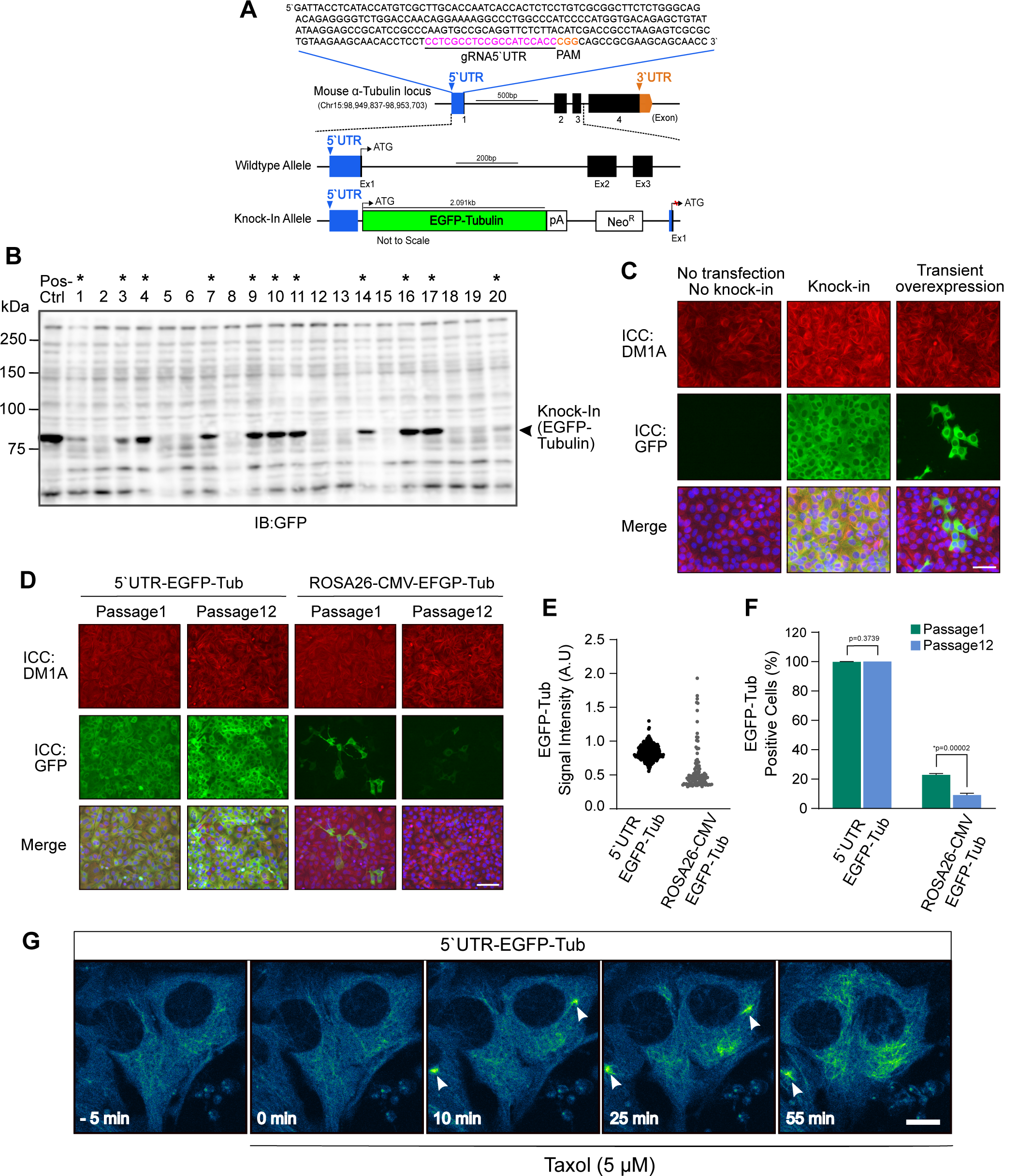
CRISPR/Cas9 mediated knock-in of EGFP-Tubulin into the 5’UTR of α-tubulin locus in IMCD3 cells. **(A)** Diagram of the mouse α-tubulin locus. The gRNA-5’UTR target sequence (magenta) upstream of the protospacer adjacent motif (PAM, orange) in the 5 untranslated region (5UTR) are indicated. **(B)** Screening of EGFP-Tubulin positive clones by western blotting. Lysate from IMCD3 cells overexpressing EGFP-Tubulin was used as positive control. Clones positive for integration are marked by asterisks*. **(C)** Fluorescence microscopy images after immunocytochemistry showing α-tubulin (DM1A; red), EGFP-Tubulin (GFP; green) and the nucleus (DAPI; blue) in control (no knock-in and no transfection of EGFP-Tubulin), knock-in cells, and cells with transient overexpression of EGFP-Tubulin (Scale bar, 10 μm). **(D)** Comparison of expression levels of endogenous EGFP-Tubulin in cells expressing EGFP-Tubulin from endogenous tubulin locus (5’UTR-EGFP-Tub) and from ROSA26 locus (ROSA26-CMV-EGFP-Tub) at passage 1 and passage 12 (Scale bar, 10 μm). **(E)** Quantification of GFP-Tubulin signals in ***D*. (F)** Quantification of EGFP-Tubulin-positive cells in ***D***. Data are shown as the mean ± s.e.m. (WT: wildtype; KI: knock-in; TF: transfection; *, *p* < 0.05). **(G)** Time-lapse confocal microscopy images of KI cells with endogenous expression of EGFP-tubulin and treated with 5 µM Taxol after 5 min into imaging. Accumulation of EGFP-Tubulin (Pseudo color: Blue and Green) could be seen in the microtubules after Taxol treatment. Arrow heads show the position of centrosomes. Time stamps are shown at the bottom (Scale bar, 10 μm).

The expression of the KI transgene was also confirmed with immunostaining. The KI EGFP-tubulin signals amplified with anti-GFP antibodies were diffused in cytoplasm, modestly overlapping with fibrous signals detected by the anti-α-tubulin monoclonal antibody in KI cells (Fig. 4C; middle row). The expression level in the KI stable cell line was more moderate and quite homogenous compared to that by the transient overexpression upon plasmid transfection (Fig. 4C; middle row versus right row). No GFP signals were detected in the negative control, i.e. neither knocked-in nor transfected, cells (Fig. 4C; left row).

To assess the long-term stability of transgene expression, the stable cell line expressing EGFP-Tubulin from 5’UTR of endogenous Tub1a locus (5’UTR-EGFP-Tubulin) and from ROSA26 locus under CMV promoter (ROSA26-CMV-EGFP-Tubulin) were sub-cultured for 3 months in the absence of G418. 5’UTR-EGFP-Tubulin showed no overt decrease in the EGFP-Tubulin expression with high homogeneity for the 3 months (Fig. 4D). In contrast, the expression of EGFP-Tubulin from ROSA26 locus looked decreased for the same time period with heterogeneity of expression levels (Fig. 4D). Quantitative analyses demonstrate the homogeneity of the modest EGFP-Tubulin expression in 5’UTR-EGFP-Tubulin knock-in cells compared to the wide range of heterogeneous expression of EGFP-Tubulin locus driven by the CMV promoter from ROSA26 (Fig. 4E). The quantitation of the EGFP-Tubulin-positive cells well shows the remarkable stability of the KI transgene in the 5’UTR of endogenous gene locus: the clone that expressed EGFP-Tubulin at the endogenous locus remained almost perfect EGFP-positive rate for 3 months while the clone that expressed EGFP-Tubulin under the CMV promoter at ROSA26 locus decreased ∼2-fold the number of EGFP-tubulin-positive cells during the 3-month culture (Fig. 4F).

We finally performed live cell imaging of microtubule formation in the 5’UTR-EGFP-Tubulin KI cells with a confocal laser microscope equipped with a sensitive detector, GaAsP-PMT. In the KI cell line, EGFP-tubulin fluorescent signals were dim and predominantly diffused in the cytoplasm before adding Taxol, a microtubule-stabilizing agent (Fig. 4G; -5 min). The time-lapse images clearly show that intense microtubule formation occurred in a few minutes around centrosomes after Taxol addition (Fig. 4F, arrow heads; Movie 2), and that microtubule networks were formed in the cytoplasm thereafter (Fig. 4F, 55 min).

## Discussion

In this study, we developed a novel strategy to achieve site-specific knock-in reporter-tagged GOI in mammalian cell lines by CRISPR/Cas9 mediated homology-independent pathway at endogenous gene loci. Using our approach of inserting tagged GOI cassette directly at 5’UTR of desired gene loci, KI efficiency as high as 80% can be achieved and is incomparable to other CRISPR/Cas9-mediated NHEJ KI strategies (Suzuki et al., 2016, He et al., 2016, Schmidt-Burgk et al., 2016, Tálas et al., 2017, Manna et al., 2019). This suggests that 5’UTR of endogenous gene loci serves as a ‘hotspot’ for gene insertion. The 20-50% of tag-negative clones could be cells that harbor the integrated donor vectors in a reverse manner, which occur in HITI-dependent repair (Suzuki et al. 2016).

One of the limitations in labeling endogenous proteins with reporter tags using NHEJ assisted KI is that the system suffers from in-del mutations at the junctions making it hard to generate exogenous transgene reporter tag and endogenous fusion genes for chimeric reporter tagged proteins by in-frame insertion (Xao et al., 2017). Our approach solves this problem by directly inserting tagged GOI cassette upstream of the coding region of endogenous loci in the 5 ‘UTR, thus simultaneously silencing the endogenous gene expression and driving the expression of KI tagged GOI from the endogenous promoter in the targeted allele. Furthermore, in our design, the donor sequences also contain eukaryotic antibiotic resistance gene cassette Neo^R^ which permits the in-frame expression of the antibiotic resistance genes for selection of KI cells, independent of the gene expression level of the GOI. The antibiotic selection marker also makes it unnecessary to use cell sorters for the selections of pure cell clones which is often costly. Once the clones are selected, there is no need to maintain the stable cell lines with selection pressure. The complete protocol, from the initial planning to the expression analysis in KI cell lines, can be executed in about six weeks. Another benefit of our approach is that the proteins can be tagged at C-terminal and N-terminal and that researchers can use mammalian expression plasmid vectors they have for their research to construct donor vectors.

In this study, we presented three examples of tagged proteins: Arl13b-Venus, Reep6-HA, and EGFP-Tubulin. We chose these proteins because they are localized in different cellular compartments, i.e. the primary cilium (Arl13b), cell membrane (Reep6), and the microtubule cytoskeleton (Tubulin). The localization patterns of the KI proteins were consistent with the shape and positions of these organelles in fixed cells. When the same proteins were expressed transiently or from GSH locus ROSA26 under the CMV promoter, strongly varying expression levels, ectopic localization, cell morphology changes as well as the loss of expression from GSH were observed. GSH loci do not entirely recapitulate the endogenous promoter in terms of spatiotemporal expression and could be silenced over time (Klatt et al., 2020). These findings demonstrate that our method can generate “truly” stable cell lines free of artifacts, correct localization with homogeneous expression, and stable insertion of tagged GOI.

The current study also highlights that stable cell lines with fluorescent reporter tagged proteins we have made are suitable for time-lapse imaging. Cytoskeleton and subcellular localizations were visualized. We were able to observe the incorporation of EGFP-Tubulin into the microtubules after Taxol treatment. This type of approach could be beneficial to study cytoskeletal dynamics. Also, we were able to see a strong and bright fluorescent signal of Arl13b-Venus in the primary cilium without any substantial photobleaching, even though we used an old confocal laser scanning microscope equipped with the “previous-generation” photomultiplier. In general, Venus has brighter fluorescent signals than EGFP but have relatively lower photostability (Shaner, Steinbach, & Tsien, 2005).

Like with any other technology there are also certain limitations to our protocol. Our method depends on the availability of the gRNA in the desired loci for integration. In eukaryotes, the median lengths of 5’UTR range from 53-218 nucleotides (Leppek et al., 2018) and it is possible that suitable gRNA recognition sites may not be available for targeted insertion because of the limited length of the region. Secondly, although NHEJ is an inherently accurate pathway, it is still prone to cause off-target integrations (Maruyama et al., 2015). The rate of such unwanted events depends on the cleavage efficiency at the on-target site, the spontaneous integration frequency of the donor, and the specificity of the gRNA that was used (Lackner et al., 2015). This situation could be improved at least in part by carefully selecting the gRNA with minimum off-target effects. It is also necessary that when interpreting the data, researchers should take into consideration that random integrations could play a role in the outcome and proper controls should be included to confirm the results.

In summary, our approach enables to drive efficient expression of GOIs under endogenous control without impacting its localization and shape within the cells and allows the generation of stable cell lines for applications such as time-lapse imaging and protein localization studies without the use of exogenous markers. This approach can easily be adapted to further fluorescent proteins or epitope tags and others. This protocol is hence a fast and adaptable methodology for establishing stable cell lines with endogenous tags such as EGFP, Venus, and HA according to the needs of the researchers.

## Acknowledgements

We thank Madoka Hamada and Rina Kamigaki for technical assistance on preparing plasmids and stable cell lines.

