## Supplementary Data for "Knock-in of labeled proteins into 5’UTR enables highly efficient generation of stable cell lines"

**Supplemental Data**

**Movie Legends**

**Movie1. Time-lapse imaging of knock-in NIH/3T3 cells primary cilium with endogenous expression of Arl13b-Venus.** Cells having a primary cilium were imaged every one minute for total of 90 min using a confocal microscope (Olympus FV1000) equipped with an oil immersion lens (60X, NA:1.35).

**Movie2. Time-lapse imaging of knock-in IMCD3 cells with endogenous expression of EGFP-Tubulin.** Cells were imaged every one minute for total of 60 min using a confocal microscope (Olympus FV3000) equipped with an oil immersion lens (60X, NA: 1.40). Cells were treated with 5µM Taxol after 5 min into imaging. Accumulation of EGFP-Tubulin (Pseudo color: Blue and Green) could be seen in the microtubules after Taxol treatment.
